# Integrin α10β1-selected mesenchymal stem cells are protective in a murine model of post-traumatic osteoarthritis

**DOI:** 10.64898/2026.08.04.742505

**Authors:** Jelena Schwarz, Xujia Wang, Marta Empere, Kristina Uvebrant, Elena Ludwig, Heidrun Grondinger, Zsuzsanna Farkas, Maximilian M Saller, Riccardo E Giunta, Evy Lundgren-Åkerlund, Attila Aszodi, Paolo Alberton

## Abstract

**Background:** Post-traumatic osteoarthritis (PT-OA) is a debilitating condition with significant unmet clinical need. Mesenchymal stem cells (MSCs) represent promising candidates for the treatment of cartilage conditions, owing to their immunomodulatory and regenerative capacities. However, the marked heterogeneity of MSC preparations remains a major challenge for product standardization and prediction of therapeutic efficacy. We previously identified integrin α10β1 as a marker for the selection of a more homogenous MSC preparation, with cartilage repair potential *in vivo*. In this study, we evaluated the therapeutic efficacy of human MSCs selected for high integrin α10β1 expression in a murine PT-OA model, and compared their effects with those of unselected MSCs.

**Methods:** Unselected or integrin α10-selected human bone marrow MSCs were characterized by flow cytometry and differentiation potential into adipogenic, osteogenic and chondrogenic lineages. Cells encapsulated into fibrin gel were applied intra-articularly at the time of surgery in the destabilization of the medial meniscus (DMM) mouse model of PT-OA. Eight weeks after DMM induction, severity of cartilage damage was assessed on Safranin O-stained sections using the OARSI scoring system. Synovitis, periarticular chondrogenesis, and osteophyte formation were evaluated histologically. OA-associated proteases, extracellular matrix degradation markers, and apoptosis were analyzed by immunohistochemistry, ELISA, and TUNEL assay. Persistence of transplanted human cells was assessed by PCR.

**Results:** Integrin α10β1-selected MSCs showed the characteristic MSCs immunophenotype and trilineage differentiation capacity. *In vivo*, treatment with integrin α10-selected MSCs significantly attenuated PT-OA-induced articular cartilage degeneration compared with both unselected MSCs and vehicle-treated controls. Further histopathological analyses revealed tendency toward reduced synovitis and periarticular chondrogenesis. Moreover, integrin α10-selected MSC treatment was associated with modest reductions in apoptotic activity and decreased expression of OA-related proteases and extracellular matrix degradation markers. Lastly, human cells were not detectable in joint tissues at the study endpoint.

**Conclusions:** Integrin α10-selected MSCs demonstrated superior chondroprotective effects compared with unselected MSCs, highlighting their potential as a standardized and efficacious cell therapy for PT-OA. These findings further validate the feasibility and safety of selection and *in vivo* administration of MSCs with high expression of integrin α10.

## Introduction

Osteoarthritis (OA) is one of the most common degenerative musculoskeletal disorders causing pain, disability and enormous socioeconomic burden worldwide. The pathogenesis of OA is multifactorial and its pathological changes interest the whole joint as an organ (1). Articular cartilage degradation, subchondral bone remodeling, osteophytes formation, ligaments and menisci degeneration, and variable degrees of synovial inflammation are typical hallmarks of OA (2). To date no successful treatment options able to cure, halt or slow the progression of the disease have been developed. Therefore, OA remains one of the main unmet clinical challenges in the orthopedic field, which has attracted considerable interest from both the public research and private biotech industry.

Mesenchymal stem cells (MSCs) are considered one of the most promising candidates for cell-based therapies of cartilage conditions owing to their ease of harvest, abundance, self-renewal capacity, multilineage differentiation potential, anti-inflammatory and immunomodulatory properties (3–5). The proclaimed potential and confident expectations from MSC therapies are reflected by the fact that among all clinical trials involving biologics for OA and cartilage defect treatments, one third of them implicates the use of MSCs. Yet, only a limited number of these trials have progressed to advanced clinical phases and only two have reached phase 4 (6). This lack of success in clinical development and finally market approval is partially due to the lack of quality control standards of MSCs used in clinical treatments. There is a substantial need for consensus on isolation, expansion and administration protocols to achieve better quality, safety and efficacy of MSCs in clinical trials (3). Furthermore, it is well-known that MSCs preparations display significant heterogeneity which is intrinsic among donors, tissue sources and cell subpopulations and that this heterogeneity can highly affect their therapeutic efficacy (7). Efforts were done towards the identification of biological markers for the isolation of potent and reproducible MSCs subpopulations specific for cartilage protection and repair, which are comprehensively reviewed in Zha et al., 2021 (8).

In this regard, we have extensively demonstrated that the collagen-binding receptor integrin α10β1 (9) represents a phenotypic marker for the identification and isolation of a homogenous MSCs preparation with enhanced chondrogenic potential, tissue homing properties and immunomodulatory potency *in vitro* (10, 11). *In vivo*, MSCs selected for integrin α10β1 expression, have shown therapeutic efficacy across several preclinical models of post-traumatic osteoarthritis (PT-OA) and cartilage injury. Specifically, these cells attenuated disease progression in an equine talar impact model of PT-OA (12), homed to a knee cartilage defect and assumed a chondrocyte-like phenotype in a rabbit chondral defect model (13), and reduced pain and cartilage degradation while enhancing immunomodulation in an equine carpal osteochondral PT-OA model (14). Furthermore, administration of integrin α10β1-selected MSCs induced broad alterations in the synovial fluid extracellular vesicle proteome (15) and modulated microRNA expression profiles (16) in experimental equine PT-OA. Collectively, these findings support integrin α10β1 as a reliable marker for the selection of potent and functionally consistent MSC populations with strong therapeutic potential for cartilage preservation and regeneration.

While the efficacy and safety of selecting and using integrin α10-expressing MSCs in *in vivo* OA animal models have been widely proved, the direct comparison with their unselected counterpart has not been investigated so far. Therefore, the aim of this study was to challenge the potential protective role of integrin α10β1-selected versus unselected MSCs in an *in vivo* murine model of PT-OA induced by the destabilization of the medial meniscus (DMM) (17).

## Materials and methods

### MSCs preparation and selection

Human MSCs were isolated from bone marrow aspirates obtained from a healthy 23-year-old male donor via density gradient centrifugation as previously described (10). Cells were cultured in DMEM/F12 (Gibco, USA) supplemented with 5% platelet lysate (Stemulate®; Cook general biotechnology, USA) and 1% penicillin/streptomycin (Gibco), under hypoxic conditions (4% O_2_, 5% CO_2_) until ∼80% confluence. MSCs were detached using Accutase (Gibco) and characterized by flow cytometry (BD Accuri™ C6, BD Biosciences) based on positive expression of CD73, CD90, and CD105, and absence of hematopoietic markers CD14, CD34, CD45, CD11b, CD19, and HLA-DR (all BD Biosciences). After expansion to passage 2, a portion of cells was labeled with a proprietary monoclonal antibody against integrin α10 (Xintela AB) and sorted for high Integrin α10-expression using fluorescence-activated cell sorting (FACSAria, BD Biosciences). This population is referred to as integrin α10β1-selected MSCs (or ITG α10β1-selected MSCs in figures). Integrin α10β1-selected MSCs were re-seeded for recovery and expansion for one additional passage before being aliquoted, frozen in Stem-Cellbanker (Amsbio, UK) and stored in liquid nitrogen until shipment on dry ice and subsequent use in *in vivo* experiments. An unsorted subset of cells from the same donor at the same passage, and not subjected to antibody labeling or sorting, served as the control cells preparation (unselected MSCs).

Three days prior to *in vivo* application, cryopreserved MSC aliquots were thawed and cultured for 48 hours. Immediately before administration, cells were detached using Accutase (Gibco), washed twice in PBS, and resuspended at 33,333 cells/µl in thrombin (Artiss kit, Baxter, USA). The cell suspension was used as described in Material and Methods section: *Surgical induction of PT-OA and MSCs application*.

### Tri-lineage differentiation

Unselected and integrin α10β1-selected MSCs were differentiated into adipogenic, osteogenic, and chondrogenic lineages as previously described (18) with minor modifications. For adipogenic differentiation, cells at 90-100% confluence were cultured for 5 days in induction medium consisting of high-glucose DMEM (Gibco) supplemented with 10% FBS, 1 mM dexamethasone, 0.2 mM indomethacin, 0.1 mg/mL insulin, and 1 mM 3-isobutyl-1-methylxanthine (all Sigma-Aldrich, USA), followed by 2 days in maintenance medium (high-glucose DMEM supplemented with 10% FBS, and 0.1 mg/mL insulin). This 7-day cycle was repeated for a total of 3 weeks. After 21 days, cells were washed with PBS, fixed with 4% PFA/PBS (Merck, Germany) for 15 min at RT, and stained with 1:2500 Bodipy staining/dH_2_O (Thermo Fischer Scientific, USA) for 15 min at 37°C. After washing, phase contrast and fluorescence images were taken with an Axiocam MRm camera mounted on an AxioObserver microscope (Carl Zeiss, Germany).

For osteogenic induction, cells at ∼80% confluency were cultured in high-glucose DMEM (Gibco) supplemented with 10% FBS, 10 mM b-glycerophosphate, 50 mM l-ascorbic acid 2-phosphate and 100 nM dexamethasone (all Sigma). Medium was replaced every 3 days for 21 days. Following stimulation, cells were washed with PBS, fixed in 4% PFA/PBS for 15 min at RT, and stained with 40 mM Alizarin Red (Sigma) for 20 minutes to assess matrix mineralization. Photomicrographs were taken with an AxioCam 105 color camera mounted on an AxioVert 40 CFL (Carl Zeiss).

For chondrogenic differentiation, 2.5×10^5^ cells/pellet were centrifuged at 500 x g for 10 min in untreated V-bottom 96-well plates (Corning, USA). Pellets were cultured for 28 days in chondrogenic medium consisting of high-glucose DMEM (Gibco) supplemented with 10 µM dexamethasone, 1 mM sodium pyruvate, 0.195 mM l-ascorbic acid, and 1% insulin transferrin selenium (all Sigma), and 10 ng/mL each of TGF-β1 and BMP-2 (R&D Systems, USA). Afterwards, pellets were fixed with 4% PFA/PBS, cryo-embedded in Tissue Tek cryomedia (Sakura, USA), cut in 7 µm slices on a cryostat Microm HM500 (Thermo Scientific, USA), and stained with Safranin O staining for the detection of sulfated glycosaminoglycans and immunostained for collagen type II (mouse monoclonal II-II6B3, 1:10, DSHB, USA) as described in the immunohistochemistry section of materials and methods. Bright-field images were taken with an AxioCam 105 color camera mounted on an AxioVert 40 CFL (Carl Zeiss).

For adipogenic and osteogenic assays, unstimulated MSCs maintained in standard growth medium served as negative controls. In the chondrogenic differentiation, control pellets were cultured in differentiation medium without growth factors. All differentiation assays were reproduced twice in triplicates.

### Surgical induction of PT-OA and MSCs application

Twelve-week-old C57BL/6j mice (Charles River, Germany) were used for the surgical induction of PT-OA. Animals were housed at the central animal facility of the LMU downtown hospital under the FELASA-guidelines, in standard conditions (12 h light/dark cycle) with *ad libitum* access to food and water. Following shipment, mice underwent a two-week acclimatization period. All animal procedures were approved by the ethics committee of the local authorities (District Government of Upper Bavaria, Germany, Animal Application: 55.2-1-54-2532-150-13). Mice were narcotized via intraperitoneal application of a mixture of Fentanyl (0,05 mg/kg KG, Fentanyl Janssen, Janssen-Cilag, Germany), Midazolam (5 mg/kg KG, Dormicum, ratiopharm) and Medetomidin (0,5 mg/kg KG, Antisedan, Orion, Finland). PT-OA was induced by surgical destabilization of the medial meniscus (DMM) as described in Glasson et al., 2007 (17). Immediately after the transection of the medial menisco-tibial ligament, MSCs were administered intra-articularly using the clinically approved fibrin sealant Artiss® (Baxter, USA) as cell carrier. Fibrinogen and thrombin components were mixed 1:1 and delivered via micro-pipette. The empty group received solely 3 µl of fibrin sealant (1,5 µl fibrinogen:1,5 µl thrombin) into the right operated knee (n=9). For the other two experimental groups, either 50.000 unselected (n=11) or integrin α10β1-selected (n=10) MSCs were firstly suspended in the 1,5 µl thrombin component and then mixed with 1,5 µl fibrinogen. Sham surgery, consisting in the solely visualization of the medial meniscotibial ligament without further intervention, served as control group (n=4). To neutralize narcosis, 0,2 ml saline solution mixed with Naloxon (1,2 mg/kg KG, Narcanti®, Bristol-Meyer, Germany), Flumazenil (0,5 mg/kg KG, Anexate®, Roche, Germany) and Atipamezol (2,5 mg/kg KG, Antisedan®, Zoetis, USA) was injected intraperitoneal. Mice were operated at 12 weeks of age and sacrificed for analysis at 8 weeks post-operation. Animals were monitored daily during the first 3 post-operative days and every third day thereafter for behavioral or mobility changes. Sacrifice was performed by cervical dislocation at study endpoint.

### Tissue processing

Right hind limbs were skinned and fixed in 4% PFA/PBS (Sigma-Aldrich, Germany), overnight at 4°C with mild agitation. Samples were decalcified for approx. 4 weeks in 20% ethylenediaminetetraacetic acid (EDTA)/PBS, pH 8.0 (Sigma-Aldrich). Following decalcification, tissues were dehydrated through graded ethanol series, cleared in xylene, and embedded in paraffin. Serial sagittal sections (6 µm) were cut using a rotary microtome (HM360, Thermo Scientific) and mounted onto Superfrost Plus glass slides (Thermo Scientific).

### Histology

General histology for Safranin O/Fast green with hematoxylin counterstaining was performed as previously described (19). For picrosirius red staining, sections were cleared in xylol 2 x 30 min, re-hydrated in a crescent series of ethanol and stained as previously described (20). Bright-field and polarized light images were acquired with an AxioVert 40 CFL using a 40x objective and AxioCam 105 color camera (Carl Zeiss).

### Scoring systems for histopathological changes of the knee joint

Articular cartilage damage following DMM surgery was evaluated by three blinded observers using the OARSI scoring system (21). The tibial and femoral cartilage were scored separately, and cumulative scores (range 0–12) combining both sites are presented. Evaluation of synovitis was performed using the score system proposed by Krenn et al. (22). The scoring system assessed three histopathological features of the synovial membrane: synovial lining layer hyperplasia, stromal cellularity, and leukocyte infiltration. Each feature was graded from 0 to 3, and the total score was calculated as the sum of the individual scores. Periarticular chondrogenesis was graded as follow: 0) normal; 1) slight; 2) moderate; 3) severe. Osteophyte formation, was graded as following: 0) no osteophyte; 1) cartilage; 2) cartilage and bone, vascular invasion; 3) bone. Synovitis, periarticular chondrogenesis and osteophyte scoring were assessed by two blind observers.

### Immunohistochemistry

Immunohistochemistry was performed as described in our previous study (19). The following primary antibodies were used: Lubricin (Abcam ab28484, 1:300), A disintegrin and metalloproteinase with thrombospondin motifs 5 (ADAMTS-5, a gift from Amanda Fosang, University of Melbourne; 1:5000), MMPs-generated aggrecan neoepitope VDIPEN (a gift from Amanda Fosang, University of Melbourne; 1:2000), matrix metollproteinase-13 (MMP-13, Sigma Aldrich MAB13424, 1:100), and types I and II collagens cleavage neoepitope generated by MMPs (C1,2C, IBEX 50-1035; 1:600). Six representative animals per DMM group and two for the Sham group, were stained and evaluated by two blinded observers.

### Enzyme-linked immunosorbent assay

In order to detect the levels of collagen type II degradation epitopes, the commercially available Mouse c-Telopeptide of Type II Collagen (CTX-II) ELISA Kit (MyBioSource MBS706197, San Diego, CA, USA) was used. Blood samples were collected from the vena cava directly after mice euthanasia and allowed to clot for 1 hour at room temperature. Afterwards, serum was collected by 15 minutes centrifugation at 1500 x g at 4°C and stored at −80°C until assayed according to manufacturer instructions.

### TUNEL assay

Apoptotic chondrocytes were detected in tissue sections using the In Situ Cell Death Detection Kit (Roche, Germany; Cat. No. 11684795910) following the manufacturer’s protocol. Slides were counterstained with DAPI-Fluoroshield™ (Sigma-Aldrich) to visualize nuclei. Fluorescence images were taken with an Axiocam MRm camera mounted on an AxioObserver microscope (Carl Zeiss). DAPI and TUNEL-positive cells were counted with the plug-in tool of Image J software (https://imagej.nih.gov/ij/,USA) and presented as percentage of TUNEL-positive cells. Six representative animals per DMM group and two for the Sham control group, were stained and analyzed.

### Detection of eventual grafted cells in knee joints

To extract genomic DNA (gDNA) from paraffin-embedded tissue sections the TaKaRa Dexpat^TM^ Easy Kit (TAKARA BIO INC., Cat. #9104, Japan) was used. Briefly, the pre-heated TaKaRa DexpatTM Easy solution was pipetted directly on tissue section and then the region of knee joint was scraped off with a scalpel. The tissue-solution mixture was transferred to microtubes and incubated at 100°C for 10 minutes, gently inverted every 2 minutes. Tubes were centrifuged at 13,000 rpm for 10 minutes at 4°C to separate the paraffin, and the aqueous phase containing genomic DNA was carefully collected and stored at 4°C. Genomic DNA was purified by adding 1/10 volume of 3 M sodium acetate followed by 2.5 volumes of ethanol, mixed by inversion, and incubated at –20°C for 1 hour. DNA pellets were collected by centrifugation at 12,000 × g for 15 minutes at 4°C, washed with 70% ethanol, air-dried, and resuspended in TE buffer for storage at 4°C. PCR was performed to detect the human-specific Charcot-Marie-Tooth Disease gene (*CMT1A*: forward 5′- gaaattcatttaaaagcattttaac-3′ and reverse 5′-gctaatagtcatgtttaaaatcatttt-3′) (Cheng et al.,), and Glyceraldehyde 3-phosphate dehydrogenase (*GAPDH*: forward 5′- caactacatggtttacatgttc-3′ and reverse 5′-gccagtggactccacgac-3′) (Böcker et al.,), was used as input control. Amplified PCR products were analyzed on a 2% agarose gel. Five representative animals per DMM group (no Sham) were analyzed.

### Statistical analysis

Statistical analysis was performed with GraphPad Prism 9.1.0 (San Diego, CA, USA). Statistical significance was tested after the determination of the Gaussian distribution and variance of results using a one-way ANOVA test. *P*-value of ≤ 0.05 was considered significant. All values represent the mean and the standard deviation. A list of the number of animals and experiment replicates is stated in the dedicated methods and figure legends.

## Results

### Integrin α10β1-selected MSCs maintain characteristic mesenchymal markers expression and tri-lineage differentiation capacity

The gating strategy for the isolation of integrin α10β1-selected MSCs is illustrated in Fig. 1A. Among viable single cells, the top 20% of integrin α10-expressing cells were sorted and collected as integrin α10β1-selected MSCs for subsequent experiments, while a fraction of the heterogeneous MSC preparation was retained as an unselected MSC control. Flow cytometric analysis demonstrated that both unselected and integrin α10β1-selected MSCs displayed a typical mesenchymal phenotype, with >99% of cells expressing the canonical MSC markers CD73, CD90, and CD105 (Fig. 1B). Conversely, the expression of hematopoietic lineage-associated markers CD34, CD11b, CD19, CD45, and HLA-DR was below 1% in both cell populations (data not shown). The frequency of integrin α10-positive cells was higher in the integrin α10β1-selected MSC population (90%) than in unselected MSCs (76%). Similarly, integrin α10 expression levels, assessed by mean fluorescence intensity (MFI), were increased in integrin α10β1-selected MSCs compared with unselected MSCs (14,678 vs. 10,470, respectively).

**Figure 1.**
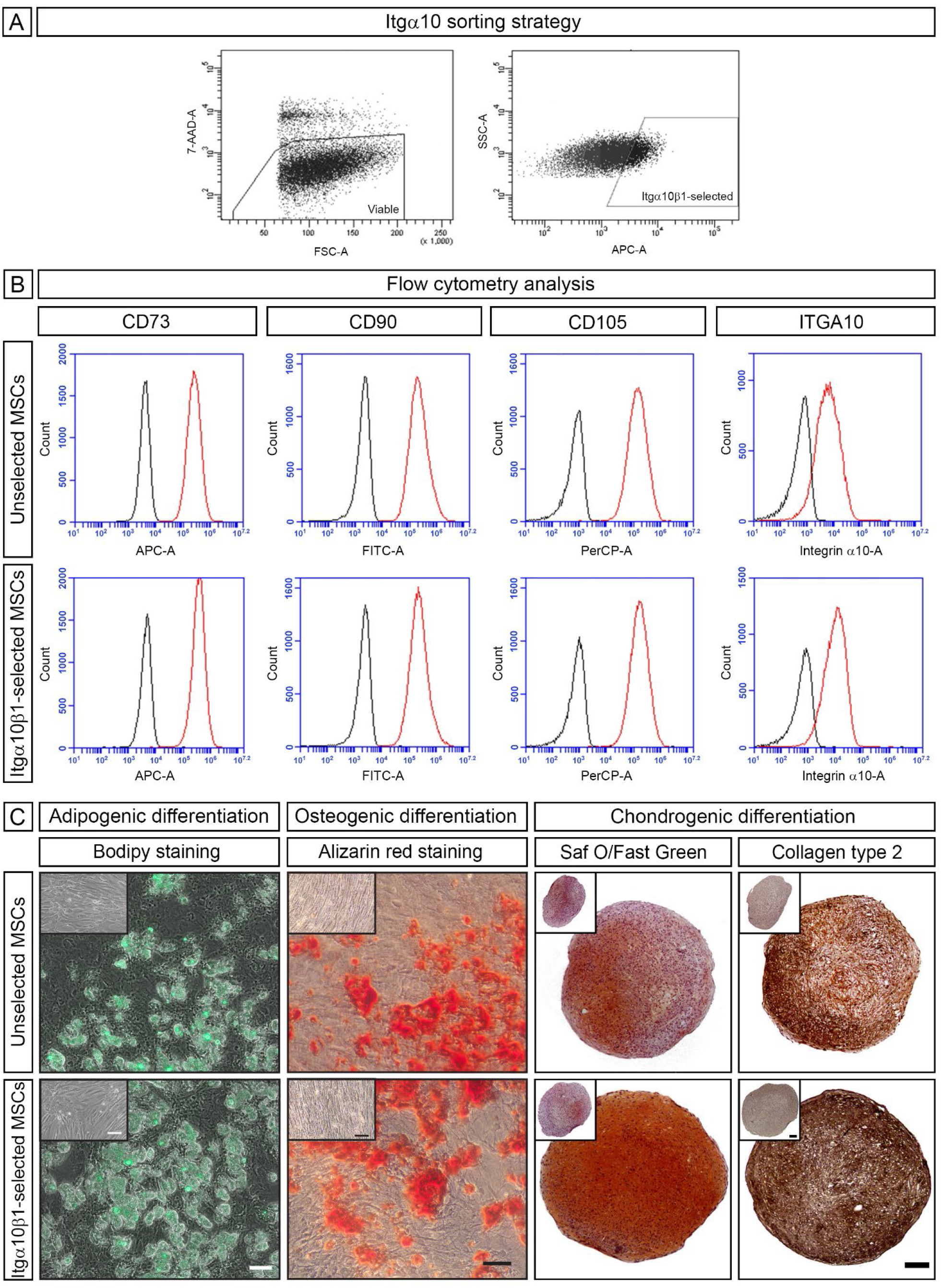
Selection and characterization of integrin α10β1-selected MSCs. (**A**) Flow cytometry dot plots illustrating gating for viable cells (left panel) and sorting strategy of MSCs with high expression of Integrin α10 (right panel), referred to as integrin α10β1-selected MSCs. Unselected MSCs from the same donor were retained as control throughout the study (unselected MSCs). (**B**) Histograms demonstrating expression of canonical mesenchymal markers CD73, 90 and 105 in both unselected and integrin α10β1-selected MSCs. The right panel shows enhanced surface expression of integrin α10 in the selected MSC population relative to unselected MSCs. (**C**) Representative images showing comparable adipogenic (Bodipy) and osteogenic (Alizarin Red) potential, but enhanced chondrogenic differentiation (Safranin O and collagen type II staining) in integrin α10β1-selected MSCs. Differentiation assays were repeated at least twice in triplicates. Scale bars: 100 µm.

The differentiation potential of MSCs, was assessed by mesenchymal tri-lineage differentiation assays. Qualitative assessment of Bodipy and Alizarin Red staining revealed comparable adipogenic and osteogenic differentiation capacities in selected and unselected MSCs (Fig. 1C). In contrast, integrin α10β1-selected MSCs exhibited enhanced chondrogenic differentiation, as demonstrated by more intense Safranin O staining and collagen type II immunoreactivity, indicative of increased sulfated glycosaminoglycan and collagen type II deposition within the chondrogenic pellets.

Altogether, these findings indicate that sorting for integrin α10 expression yields a MSC preparation that preserves typical MSC marker expression and mesenchymal differentiation potential, with superior chondrogenic capacity.

### Integrin α10β1-selected MSCs are chondroprotective in a surgical model of PT-OA

To evaluate cartilage degradation following MSCs treatment in the DMM model, histopathological assessment was performed using the OARSI scoring system (21) (Fig. 2). Successful induction of PT-OA was confirmed by significant increase in mean OARSI scores across all DMM-operated groups (empty, unselected MSCs, and α10β1-selected MSCs) relative to sham controls. Notably, while both the empty hydrogel and unselected MSC-treated groups exhibited significantly elevated OARSI scores compared to sham (\*\*\*\**p* < 0.0001), the α10β1-selected MSC group demonstrated the lowest mean scores compared with the sham group (\*\**p* = 0.0054). The empty and unselected MSC groups showed comparable cartilage damage (5.028 ± 1.189 vs. 5.023 ± 1.186, respectively), whereas treatment with α10β1-selected MSCs resulted in a significantly lower mean OARSI score (3.425 ± 1.564), differing from both the empty (\**p* = 0.0475) and unselected MSC (\**p* = 0.0351) groups. Collectively, these findings indicate that α10β1-selected MSCs confer chondroprotective effects and mitigate structural cartilage deterioration following PT-OA induction.

**Figure 2.**
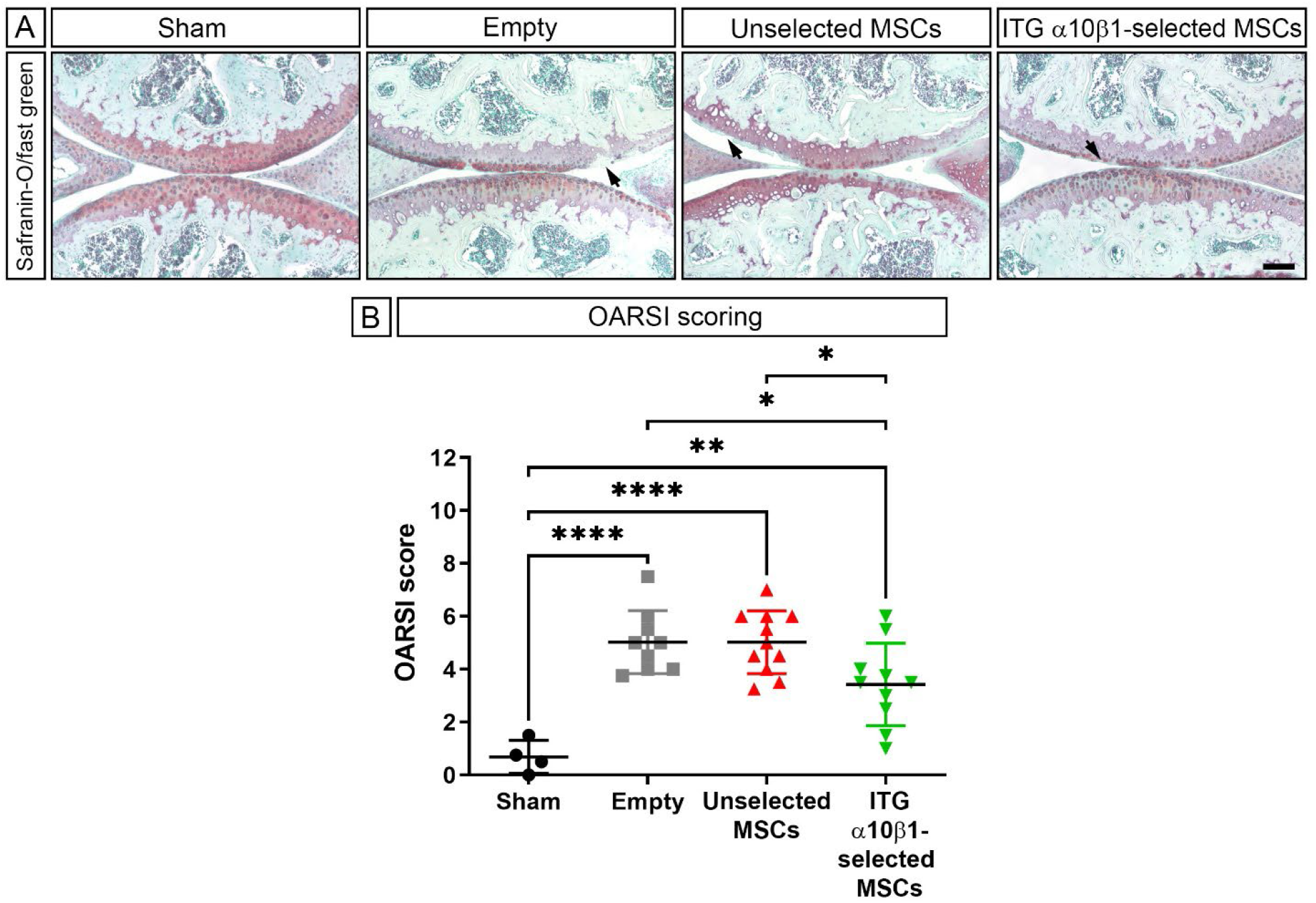
Integrin α10β1-selected MSCs attenuate cartilage degeneration in PT-OA. (**A**) Representative images of Safranin O/Fast green stained sections showing typical features of the average cartilage damage score for each group (black arrows). Scale bar: 100 µm. (**B**) Evaluation of cartilage structural damage using the OARSI scoring system. Each data point represents the average score (tibia + femur) assessed independently by three blinded observers. Data are presented as the mean ± SD. Statistical significance calculated by one-way ANOVA where \**p* < 0.5; \*\**p* < 0.01; \*\*\*\**p* < 0.0001.

Next, we examined the distribution of lubricin, a proteoglycan with essential boundary lubricant function in articular cartilage. Lubricin immunostaining was observed in the pericellular matrix of chondrocytes across the superficial and middle zones in all groups at comparable levels. However, surface-associated lubricin (black arrows, Fig. 3A) was more frequently detected in sham and integrin α10β1-selected MSC-treated joints than in the empty and unselected MSC groups. In addition, picrosirius red staining under polarized light revealed comparable collagen network organization in the deep and middle zones across all groups. By contrast, the superficial zone of sham and integrin α10β1-selected MSC-treated mice exhibited stronger and more continuous birefringence (white arrows, Fig. 3B), consistent with preservation of collagen fibrillar integrity. In the empty hydrogel and unselected MSC groups, birefringence was reduced or discontinuous, indicating superficial collagen disruption.

**Figure 3.**
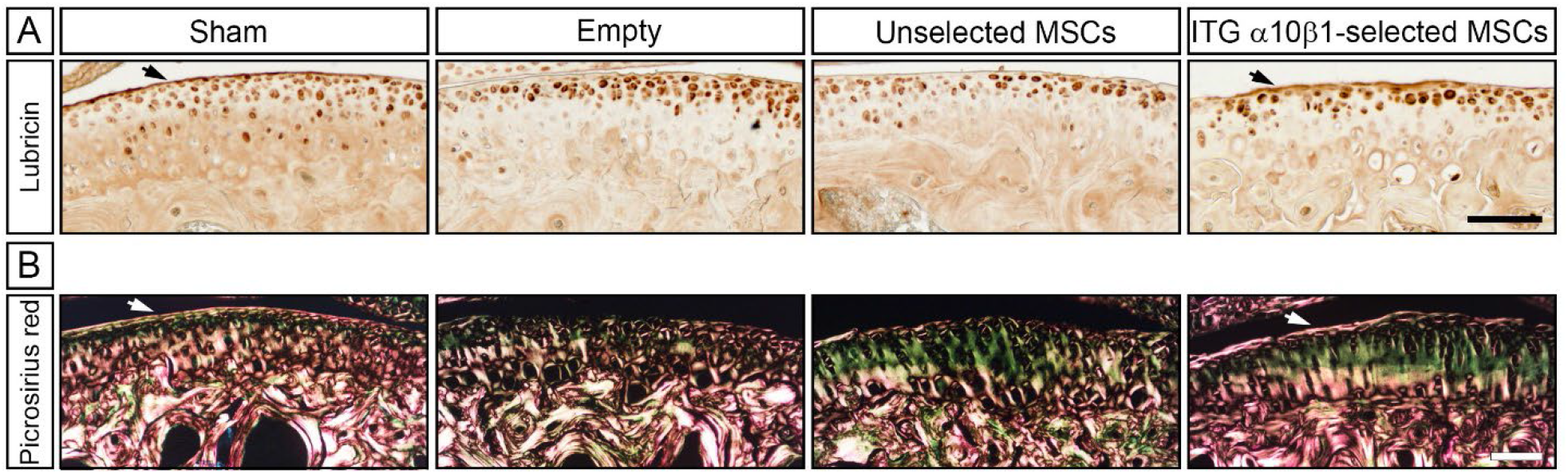
Enhanced lubricin deposition and preserved collagen architecture in joints treated with integrin α10β1-selected MSCs. (**A**) Immunohistochemical staining for lubricin (brown signal) in the pericellular matrix of non-calcified cartilage chondrocytes. Strong lubricin expression is observed in the superficial zone of sham and integrin α10β1-selected MSC-treated joints (black arrows). (**B**) Picrosirius red staining under polarized light demonstrates a continuous, well-organized collagen network at the articular surface in the sham and integrin α10β1-selected MSCs groups (white arrows). Scale bar: 100 µm.

As OA is a whole-joint disease, additional histopathological features were evaluated, including synovial pathology, periarticular chondrogenesis, and osteophyte formation (Fig. 4A–C). Synovial abnormalities were assessed using a composite score based on pannus formation, synovial membrane thickening, and sub-synovial hyperplasia. The synovitis scores (Fig. 4A) of the empty (2.67 ± 1.17) and unselected MSCs (2.32 ± 1.33) groups showed the highest deviation compared to the sham group (0.75 ± 0.65). Treatment with integrin α10β1-selected MSCs resulted in a slightly reduced synovitis score (2.00 ± 1.16) when compared to the other two DMM operated groups, however this did not reach any statistical significance. Similarly, periarticular chondrogenesis scores (Fig. 4B) were numerically lower in the α10β1-selected MSC group (0.70 ± 0.63) than in the unselected MSC (1.32 ± 0.81) and empty (1.11 ± 0.74) groups, but again without statistical significance. Sham animals displayed minimal ectopic cartilage formation (0.50 ± 1.00), except for one outlier. Osteophyte formation (Fig. 4C) showed no marked differences between the MSC-treated groups (unselected MSCs: 1.14 ± 1.33; integrin α10β1-selected MSCs: 1.10 ± 1.13), while the empty group exhibited marginally lower mean score (0.67 ± 0.83) compared with the MSCs-treated DMM groups. As with the other histopathologic features, the mean osteophyte score was minimal in the sham group (0.25 ± 0.29). In summary, high inter-individual variability across all three parameters—synovitis, periarticular chondrogenesis, and osteophyte formation—precluded the detection of statistically significant effects of MSC treatment on these secondary OA features.

**Figure 4.**
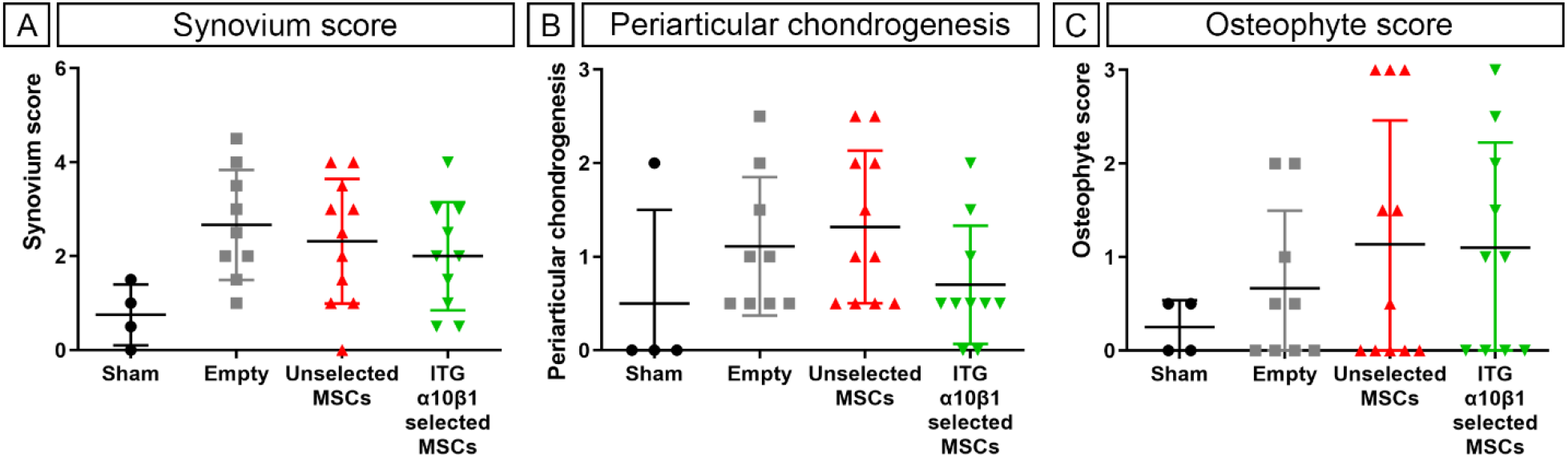
Histopathological scoring of OA-related joint changes shows no significant differences among the DMM-operated groups. Quantification of (**A**) synovial abnormalities, (**B**) periarticular chondrogenesis, and (**C**) osteophyte formation. Data represent mean scores ± SD from two blinded observers. One-way ANOVA revealed no significant group differences.

### Administration of integrin α10β1-selected MSCs is associated with reduced expression of cartilage catabolic markers

To explore potential mechanisms underlying the chondroprotective effects of MSC treatment, the expression of key mediators of extracellular matrix (ECM) degradation was evaluated by immunohistochemistry (Fig. 5). MMP-13, a major collagenase implicated in OA pathogenesis, was expressed at low levels across all groups but appeared more prominent in the empty and unselected MSC-treated joints, particularly in areas of cartilage degeneration (black arrows, MMP-13 panel). Similarly, immunoreactivity for the C1,2C neoepitope, a marker of type II collagen cleavage, was minimal in sham-operated and integrin α10β1-selected MSC-treated joints but increased in the empty and unselected MSC-treated groups. A comparable pattern was observed for ADAMTS-5, the principal aggrecanase responsible for cartilage aggrecan degradation in mice, and for the aggrecan degradation neoepitope VDIPEN. Both markers exhibited slightly increased staining intensity in the empty group compared to sham-operated controls and MSCs-treated joints. Notably, integrin α10β1-selected MSC-treated animals consistently displayed lower immunoreactivity for these catabolic markers than animals receiving unselected MSCs. Collectively, these findings suggest that treatment with integrin α10β1-selected MSCs may attenuate cartilage ECM catabolism. Although the observed differences were modest and primarily qualitative, the results are consistent with the reduced cartilage degeneration observed in this treatment group.

**Figure 5.**
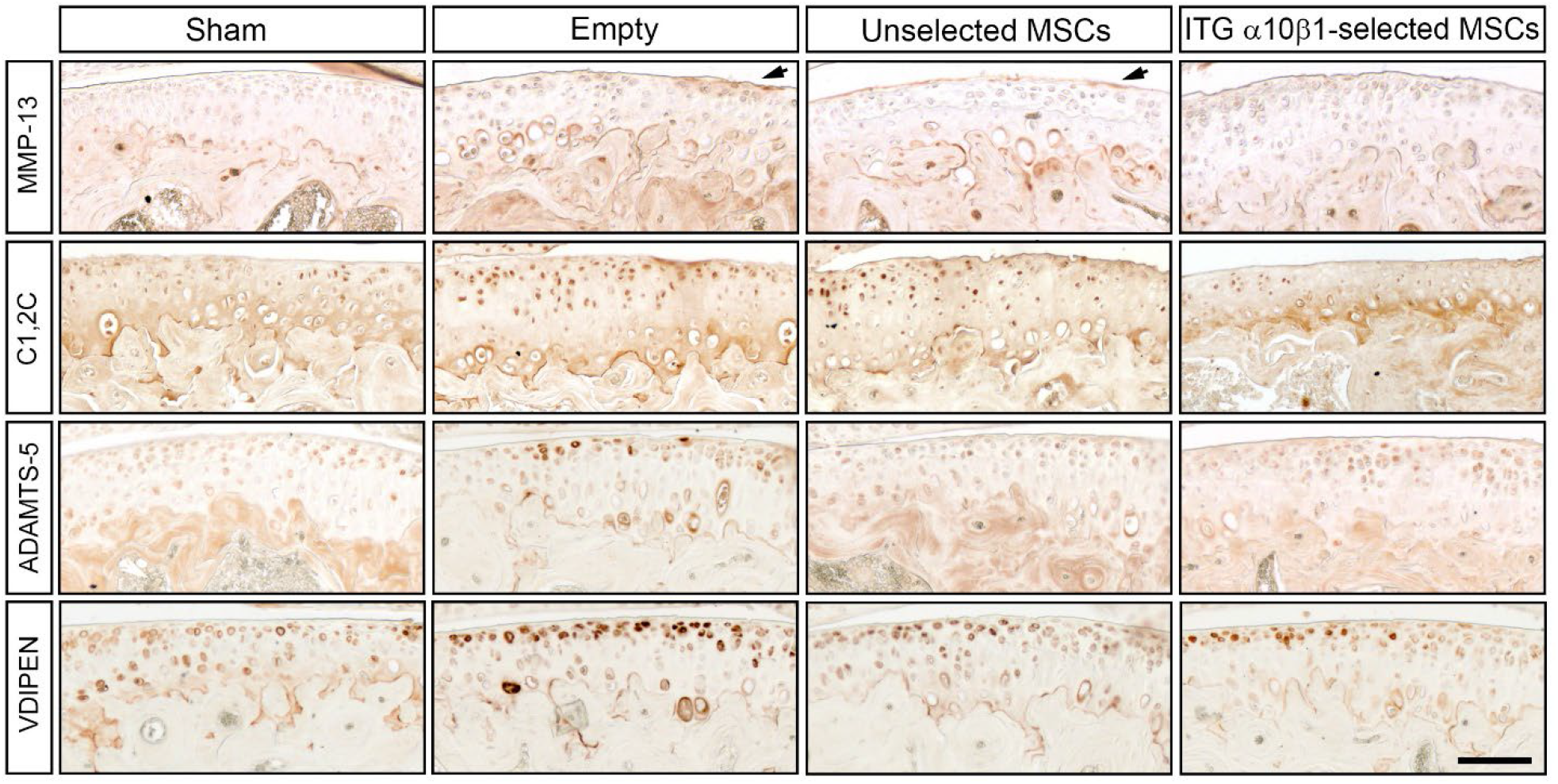
Integrin α10β1-selected MSCs slightly decrease expression of catabolic markers. Representative immunohistochemistry images showing the expression of MMP-13, C1,2C Collagen type II degradation neoepitope, the aggrecanase ADAMTS-5 and the VDIPEN Aggrecan degradation neoepitope. Scale bar: 100µm.

**Figure 6.**
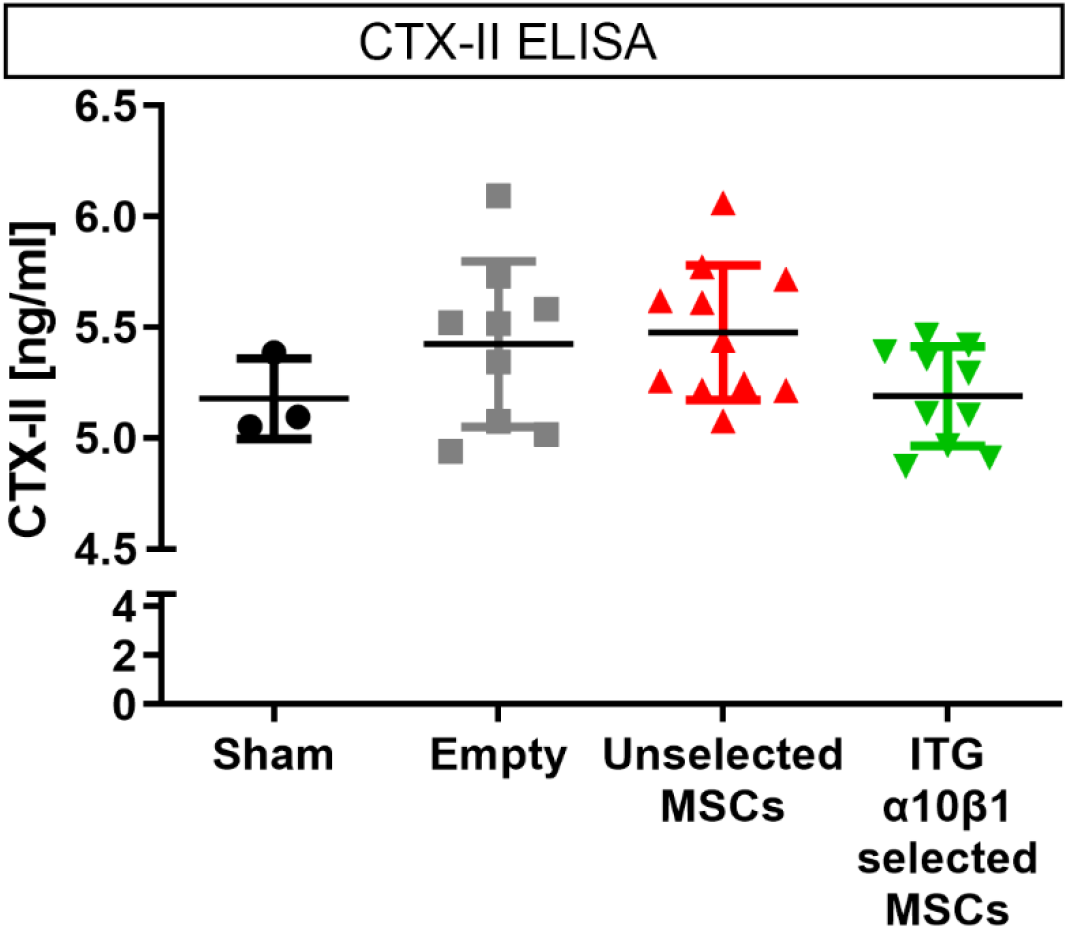
Comparable serum CTX-II levels in mice treated with integrin α10β1-selected MSCs. Scatter plot showing individual serum CTX-II concentrations with group means ± SD. One sample from the sham group was excluded because utterly out of the standard curve and therefore not considered as reliable. One-way ANOVA revealed no significant differences among the groups.

To further assess cartilage degradation, serum levels of C-telopeptide fragments of type II collagen (CTX-II) were measured 8 weeks post-surgery. Consistent with the reduced structural cartilage damage observed histologically (Fig. 2) and the lower C1,2C immunostaining detected in the integrin α10β1-selected MSC-treated group (Fig. 5), this group exhibited the lowest CTX-II levels (5.189 ± 0.223 ng/ml), closely resembling that of the sham group (5.178 ± 0.181 ng/ml). Despite the clear trend and values distribution between the empty (5.425 ± 0.373 ng/ml) and unselected (5.476 ± 0.304 ng/ml) groups compared to the sham and integrin α10β1-selected MSCs group, no statistical significance was reached.

Articular cartilage homeostasis is critically dependent on the viability of resident chondrocytes. To evaluate chondrocyte apoptosis, TUNEL staining was performed on cartilage sections (Fig. 7A). Quantification of TUNEL-positive cells (Fig. 7B) showed the lowest proportion of apoptotic chondrocytes in the integrin α10β1-selected MSC-treated group (7.51% ± 3.80), whereas the empty DMM group exhibited the highest level of apoptosis (9.50% ± 3.95). Intermediate values were observed in the unselected MSC-treated (8.75% ± 1.93) and sham-operated (8.30% ± 1.02) groups. Although these differences did not reach statistical significance and were accompanied by substantial inter-animal variability, the findings indicate a trend toward reduced chondrocyte apoptosis following treatment with integrin α10β1-selected MSCs, suggestive of a potential chondroprotective effect.

**Figure 7.**
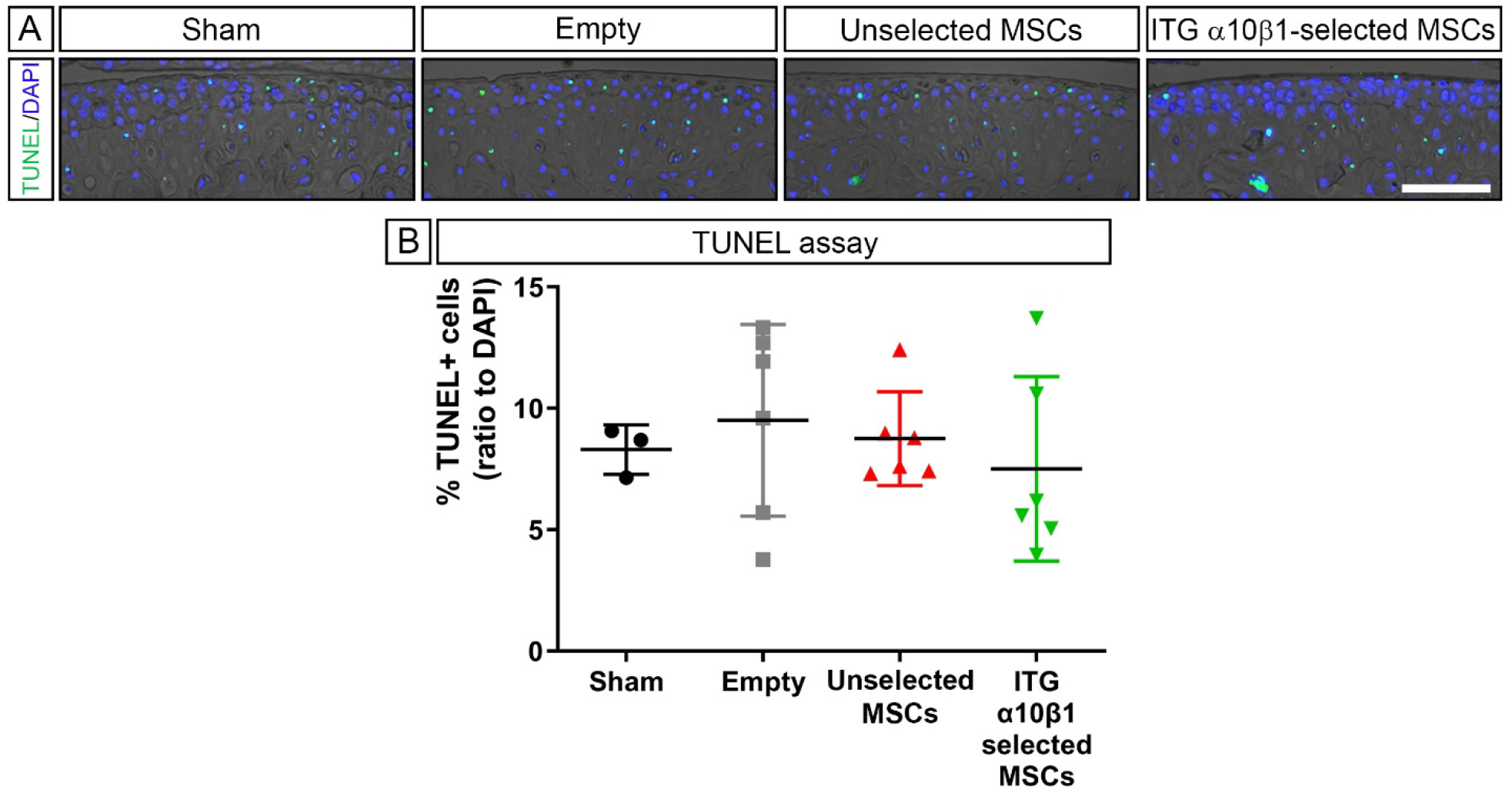
Integrin α10β1-selected MSCs show a trend toward reduced chondrocyte apoptosis. (**A**) Representative images of TUNEL staining in articular cartilage from sham and DMM groups. Phase contrast; Green: TUNEL; Blue: DAPI. Scale bar: 100 µm. (**B**) Quantification of TUNEL positive cells as a percentage of total chondrocytes in the articular cartilage. Data are presented as mean ± SD. Statistical analysis was performed using one-way ANOVA; no statistically significant differences were observed between groups.

### Human MSCs were not detectable in mouse joints eight weeks after transplantation

To evaluate the persistence and potential engraftment of transplanted human MSCs, PCR analysis targeting the human-specific CMT1A gene was performed on genomic DNA extracted from paraffin-embedded knee joint tissues. Assay sensitivity was first established using mixed mouse/human cell preparations, demonstrating a detection threshold between 500 and 1,000 human cells per 100 ng/µL genomic DNA input (Fig. 8A). Subsequent analysis of experimental samples, using 200 ng/µL genomic DNA per reaction, revealed no detectable CMT1A amplification in any treated animal (Fig. 8B). These findings indicate that transplanted human MSCs were either absent or present below the assay’s detection limit at 8 weeks post-transplantation.

**Figure 8.**
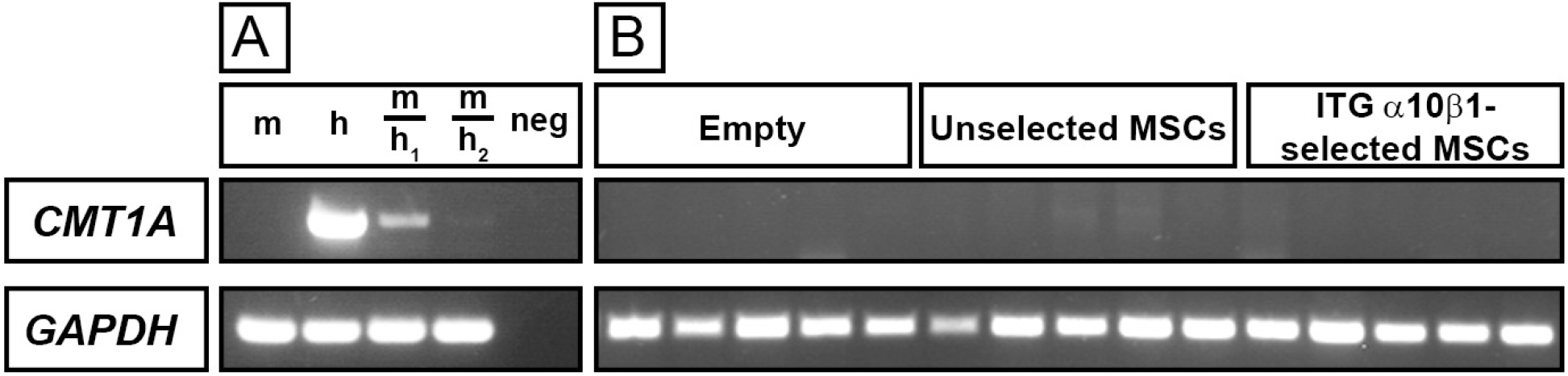
Absence of detectable human MSCs in murine knee joints 8 weeks after intra-articular administration. (**A**) PCR assay sensitivity for detection of the human CMT1A gene (345 bp). Lanes: m, 100 ng/µL genomic DNA (gDNA) from 1 × 10⁶ mouse fibroblasts; h, 100 ng/µL gDNA from 5 × 10⁴ human MSCs; m/h_1_, 100 ng/µL gDNA from a mixture of 1 × 10⁶ mouse fibroblasts and 1 × 10³ human MSCs; m/h_2_, 100 ng/µL gDNA from a mixture of 1 × 10⁶ mouse fibroblasts and 5 × 10² human MSCs. (**B**) PCR screening for CMT1A in gDNA isolated from knee joint tissues of five representative DMM-operated mice per treatment group. No human MSCs-derived signal was detected at 8 weeks post-administration. Amplification of GAPDH (181 bp) served as a DNA input and quality control. Genomic DNA input was 200 ng/µL per reaction.

## Discussion

The present study demonstrates that intra-articular administration of human integrin α10β1-selected MSCs in fibrin sealant significantly attenuated the development of PT-OA in the murine DMM model. Compared with unselected MSCs, integrin α10β1-selected MSCs provided superior chondroprotection, as demonstrated by reduced cartilage degeneration, preservation of cartilage homeostasis, and decreased collagen degradation. These findings support integrin α10β1-based selection as a strategy to improve the consistency and therapeutic efficacy of MSC-based therapies for PT-OA.

MSCs have long been considered promising candidates for regenerative medicine, including cartilage conditions, because of their regenerative and immunomodulatory properties. Nevertheless, their clinical translation has been limited by inconsistent therapeutic outcomes, largely attributed to the inherent heterogeneity of MSC preparations (7). We previously identified integrin α10β1 as a marker that enables the isolation of a homogeneous MSC preparation with enhanced potency and functional consistency (10, 11). In this study, we challenged the therapeutic potential of human MSCs selected for high expression of integrin α10β1 in a murine model of PT-OA, demonstrating their improved ability to prevent typical PT-OA-related pathological changes, when compared to their unselected counterpart or empty group controls.

The most striking finding of this study was the significant reduction in articular cartilage degeneration following treatment with integrin α10β1-selected MSCs. Although differences in synovitis, ectopic chondrogenesis and osteophyte formation were not statistically significant, a tendency for lower synovitis and periarticular chondrogenesis was nevertheless observed, consistent with the multifactorial nature of OA progression.

Preservation of cartilage structure was accompanied by increased lubricin expression in joints treated with integrin α10β1-selected MSCs. Consistent with these findings, our previous study using an equine talar impact model also demonstrated more extensive lubricin immunostaining at the articular surface and within superficial and middle zone chondrocytes in MSC-treated joints compared with untreated controls (12). Loss of functional lubricin in human and mice results in elevated coefficient of friction and consequent articular cartilage structural damage (23, 24), and several studies have demonstrated that intraarticular injection of native or recombinant lubricin can prevent cartilage degeneration and promote chondroprotection (25–28). The increased lubricin expression observed in the present study therefore indicates improved maintenance of articular cartilage surface homeostasis. Whether this reflects preservation of the superficial cartilage layer, thereby limiting lubricin loss, or enhanced lubricin production by resident chondrocytes in response to MSC-derived signals remains to be determined.

Maintenance of cartilage homeostasis was further supported by reduced expression of cartilage degradation markers. PT-OA is characterized by a progressive imbalance between anabolic and catabolic processes, leading to extracellular matrix breakdown. Among the enzymes involved, MMP-13 is a major mediator of type II collagen degradation (29), and its increased expression has been linked to cartilage destruction in human OA (30). Moreover, MMP13-overexpressing mice develop spontaneous OA-like pathology (31), whereas MMP13 deficiency confers protection against cartilage degeneration (32). In the present study, integrin α10β1-selected MSC treatment reduced both MMP13 and the collagen degradation epitope C1,2C compared with unselected MSCs and untreated controls, indicating reduced collagen breakdown and preservation of the cartilage collagen network. In contrast, expression of ADAMTS-5 and the aggrecan degradation neo-epitope VDIPEN was not significantly altered between experimental groups. These findings may suggest that, the protective effects of integrin α10β1-selected MSCs are associated predominantly with preservation of the collagen network rather than inhibition of aggrecan degradation. Together with the increased lubricin expression and reduced cartilage degeneration, these data indicate that integrin α10β1-selected MSCs promote maintenance of cartilage structural integrity through complementary mechanisms that preserve both the articular surface and ECM.

The preservation of cartilage integrity observed following treatment with integrin α10β1-selected MSCs was accompanied by a modest, although not statistically significant, reduction in chondrocyte apoptosis. Chondrocytes are the sole resident cells responsible for maintaining cartilage ECM homeostasis. During the course of OA, chondrocytes death *via* apoptosis or necrosis occurs at a higher rate than in healthy cartilage (33–35). It is long accepted that chondrocyte death and articular cartilage degradation are closely linked, as regions of increased apoptosis colocalize with proteoglycan depletion (36), while collagen network disruption can itself trigger apoptosis (37). Moreover, pharmacological inhibition of caspases, key effectors of apoptosis, attenuated proteoglycan loss and cartilage lesion severity in both *in vitro* (38) and *in vivo* (39) OA models, further supporting a causal link between apoptosis and cartilage degeneration. Although the reduction in apoptosis observed here did not reach statistical significance, it paralleled the lower OARSI scores and increased lubricin expression in joints treated with integrin α10β1-selected MSCs. Beyond its well-established role in boundary lubrication, lubricin protects chondrocytes from apoptosis induced by excessive mechanical loading and shear stress (40). Furthermore, although *in vivo* evidence remains limited, several *in vitro* studies have shown that MSCs-derived secretomes and extracellular vesicles exert anti-apoptotic effects on OA chondrocytes. Coculture experiments demonstrated that the adipose-derived MSC secretome protects OA chondrocytes from induced apoptosis (41). Follow-up studies showed that murine bone marrow MSC-derived extracellular vesicles also convey anti-apoptotic signals to OA-like chondrocytes (42). Similarly, human MSC-derived exosomes were shown to inhibit IL-1β-induced apoptosis in murine chondrocytes via the miR-206/GRKIP1 axis (43). Together, these observations suggest that integrin α10β1-selected MSCs may contribute to cartilage preservation, at least in part, by supporting chondrocyte survival. Whether this reflects a direct effect of MSC-derived factors or is secondary to improved preservation of cartilage structure remains to be established.

The mechanisms by which MSCs mediate tissue repair remain the subject of ongoing investigation. Although their multilineage differentiation potential initially suggested that cartilage regeneration relied on engraftment and direct tissue replacement, accumulating evidence indicates that their therapeutic effects are mediated predominantly through transient paracrine signaling rather than long-term persistence within the joint. Especially based on studies with unselected MSC preparations, long-term engraftment is often the exception rather than the rule. In the present study, no transplanted human MSCs were detected eight weeks after intra-articular administration. This finding contrasts with our previous studies in larger animal models, in which allogeneic integrin α10-selected equine MSCs remained detectable within osteochondral repair tissue 52 days following intra-articular administration in an equine PTOA model (14). Similarly, in a rabbit cartilage defect model, human integrin α10-selected MSCs demonstrated homing and engraftment within the repair tissue, accompanied by chondrogenic differentiation (13). The absence of detectable human MSCs in the present study is likely attributable to several methodological and biological factors. First, differences in the mode of cell delivery may have influenced cellular biodistribution and retention. In the current study, MSCs were encapsulated within a fibrin hydrogel prior to intra-articular administration, a strategy that may have limited cell migration and homing to sites of tissue injury. In contrast, in our previous studies, integrin α10β1-selected MSCs were administered as freely suspended cells in cryopreservation medium (13, 14), thereby facilitating their migration, engraftment, and subsequent localization within the repair tissue.

Second, the experimental models differ substantially in their biological context. In the previously described rabbit cartilage defect model (13), MSCs were delivered following the creation of a focal osteochondral defect, which might provide a local chemotactic and regenerative microenvironment that is likely to promote cell recruitment, engraftment, and differentiation. Additional factors may also have contributed to the lack of detectable transplanted cells, including the substantially smaller murine joint cavity, accelerated tissue turnover, and the lower absolute number of administered MSCs relative to our previous large-animal studies. Collectively, these factors may have reduced cell persistence below the detection threshold of our analyses. Nevertheless, these findings are consistent with the prevailing paradigm that durable MSC engraftment is not a prerequisite for therapeutic efficacy and that clinical benefit can persist despite rapid clearance of transplanted cells. The inability to detect transplanted cells at the experimental endpoint also represents a limitation of the present study. Assessment of earlier post-injection time points would have provided valuable information regarding cell retention and persistence within the joint and should be incorporated into future investigations. Likewise, the relative contribution of soluble mediators and extracellular vesicle-associated cargo to the observed therapeutic effects remains to be determined.

The superior *in vivo* efficacy of integrin α10β1-selected MSCs is likely attributable to their enhanced biological properties. Before transplantation, we analysed the characteristics of the two MSCs preparations. Integrin α10β1-selected cells demonstrated the typical MSC immunophenotype and trilineage differentiation capacity while exhibiting enhanced chondrogenic potential, consistent with previous reports in human bone marrow-derived MSCs (10) and equine adipose-derived MSCs (11). These findings further support integrin α10β1 as a marker for isolating a functionally distinct and more homogeneous MSC subpopulation with superior chondrogenic capacity.

The therapeutic efficacy of MSCs is strongly influenced by the experimental context, including the disease model, timing of administration and delivery strategy. To evaluate the effects of integrin α10β1-selected MSCs, we employed the destabilization of the medial meniscus (DMM) model, a well-established and reproducible model that closely recapitulates the progressive structural changes observed in human PT-OA (17, 44). The 8-week endpoint was selected to assess established OA while avoiding end-stage cartilage destruction that could mask eventual therapeutic effects.

The timing of MSC administration is recognized as a critical determinant of therapeutic outcome. Previous studies have shown that early delivery of adipose-derived MSCs attenuates inflammation and joint pathology in collagenase-induced OA, whereas delayed treatment is less effective (45). Likewise, MSC administration following DMM induction has yielded limited protection despite efficacy in the more inflammatory collagenase model (46), suggesting that MSCs may exert their greatest benefit by modulating the early inflammatory response after joint injury. Consistent with this concept, in our study, a single intra-articular injection of encapsulated integrin α10β1-selected MSCs at the time of surgery significantly attenuated PT-OA progression and provided greater chondroprotection than unselected MSCs. Together, these findings support integrin α10β1-based selection as an effective strategy to enhance the disease-modifying potential of MSC therapy.

In this study, we investigated fibrin hydrogel as delivery vehicle to support the MSCs therapy. Fibrin hydrogels are widely used for MSC encapsulation because they support cell survival, adhesion and homogeneous distribution while providing a biodegradable matrix that can be generated from autologous sources (47–49). As a provisional ECM rich in integrin-binding motifs, fibrin further promotes cell retention and viability following transplantation (50, 51). Its effectiveness as a carrier has been demonstrated in several regenerative applications, including cartilage repair, where fibrin-based delivery improves cell survival and tissue regeneration compared with cell administration alone (50, 52–54). These characteristics support its use in the present study. Nevertheless, the absence of a cell-only treatment group represents a limitation, as fibrin itself may have contributed to joint protection by improving the local biomechanical environment. Inclusion of such a control in future studies would allow the independent contribution of the carrier to be more accurately defined.

## Conclusions

This study provides further evidence that selecting MSCs based on high surface expression of integrin α10 identifies a homogenous, consistent, functionally distinct, and therapeutically potent cell preparation with enhanced capacity to protect and preserve articular cartilage. Notably, our findings suggest that integrin α10β1-selected MSCs may represent a promising strategy for early intervention following traumatic joint injury. Importantly, this is the first *in vivo* study demonstrating of the superior efficacy of integrin α10β1-selected MSCs compared with unselected, heterogeneous MSC preparations in mitigating cartilage degeneration associated with PT-OA. In summary, this study further supports the therapeutic potential of integrin α10β1-selected MSCs as a targeted cell-based approach for the treatment of cartilage conditions.

## Abbreviations

OA: Osteoarthritis
MSCs: Mesenchymal stem cells
PT-OA: Post-traumatic osteoarthritis
DMM: Destabilization of the medial meniscus
ECM: Extracellular matrix

## Aknowledgements

The authors would like to thank Dr. Lucienne Wonk for excellent review of the manuscript.

## Author contributions

Conceptualization: E.L-A., K.U., A.A., P.A.; methodology: J.S., M.E, X.W., H.G., K.U., Z.F., M.M.S., A.A, P.A; data analysis: J.S., K.U, M.E, X.W., M.M.S. E.L-A., A.A., P.A.; writing-original draft preparation: J.S, A.A., P.A.; writing-review and editing: M.M.S., R.E.G., E.L-A., K.U., A.A, P.A.; funding acquisition: E.L-A., A.A.. All authors have read and agreed to the published version of the manuscript.

## Availability of data and materials

All relevant data is contained within the manuscript. Additional data can be provided upon reasonable request.

## Declarations

### Ethics approval and consent to participate

All animal experiments were approved by the ethics committee of the local authorities (District Government of Upper Bavaria, Germany, Animal Application: 55.2-1-54-2532-150-13).

### Consent for publication

Not applicable.

### Author details

^1^ Musculoskeletal University Center Munich (MUM), Department of Orthopaedics and Trauma Surgery, Ludwig-Maximilians-University (LMU), Munich, Germany; ^2^ Xintela AB, Medicon Village, 223 81 Lund, Sweden; ^3^ Division of Hand, Plastic and Aesthetic Surgery, LMU University Hospital, LMU Munich

